# Wave propagation enhances extracellular signal strength in small excitable tissues

**DOI:** 10.1101/2025.07.07.663416

**Authors:** Karoline Horgmo Jæger, Aslak Tveito

## Abstract

Human induced pluripotent stem cell-derived cardiomyocytes (hiPSC-CMs) offer new opportunities to study cardiac dynamics and drug responses. High-throughput platforms often rely on extracellular potential (EP) recordings as a proxy for transmembrane voltage, making EP signal strength critical for analysis. EPs arise from spatial inhomogeneities – either intrinsic or induced by wave propagation. Using a computational model, we show that uniform simultaneous stimulation of all cells suppresses wave propagation and results in weak signals. In contrast, localized stimulation that initiates a traveling wave generates much stronger EPs, improving the robustness of biomarker extraction from extracellular recordings.

## 1 Introduction

The development of cardiomyocytes derived from human-induced pluripotent stem cells (hiPSC-CMs) has enabled *in vitro* studies of human cardiac cell behavior [1, 2, 3]. These cells are increasingly used to assess disease mechanisms and drug-induced effects [4, 5, 6, 7]. To support large-scale testing, hiPSC-CMs are commonly cultured in multiwell platforms or integrated into microphysiological systems (MPSs) [8, 9, 10]. These platforms often rely on extracellular potential (EP) recordings to infer electrophysiological activity [11, 12, 13]. Here we show that the interpretability of EPs depends critically on the stimulation strategy.

It was shown in [14] that autonomous, fully homogeneous cells may generate no measurable EP, even during full action potentials. The same holds for networks of identical, synchronously firing cells. To analyze such systems using extracellular signals, spatial voltage gradients must be introduced – typically by applying external stimulation. These gradients give rise to measurable EPs. Here, we examine how different stimulation protocols shape the spatial activation pattern and thereby influence the extracellular signal.

Using a computational model, we compare four conditions: (i) no stimulation, where cells beat autonomously; (ii) weak uniform global stimulation; (iii) strong uniform global stimulation; and (iv) localized stimulation to initiate wave propagation. Our computations show that conditions (i)–(iii) yield weak EPs, whereas condition (iv) produces strong, well-defined signals due to the traveling excitation wave. This demonstrates that the stimulation protocol plays a critical role in generating measurable extracellular signals.

## 2 Methods

### 2.1 Computational model

We perform simulations of a collection of hiPSC-CMs using the Kirchhoff network model (KNM) [15]. The model parameters, cell collection geometry, and the model for the single-cell dynamics are adapted from Chip C1 in [16]. The cell collection consists of a two dimensional sheet of 1637 hiPSC-CMs connected to their neighbors by a default gap junction strength of *G*_*g*_ = 1.7 nS. The cell collection is surrounded by an extracellular bath, and Dirichlet boundary conditions of the form *u*_*e*_ = 0 mV are applied on the outer boundary of this bath. In Figure 1A, the geometry of the cell collection is illustrated. In addition, the location of stimulation and measurement electrodes are indicated.

**Figure 1:**
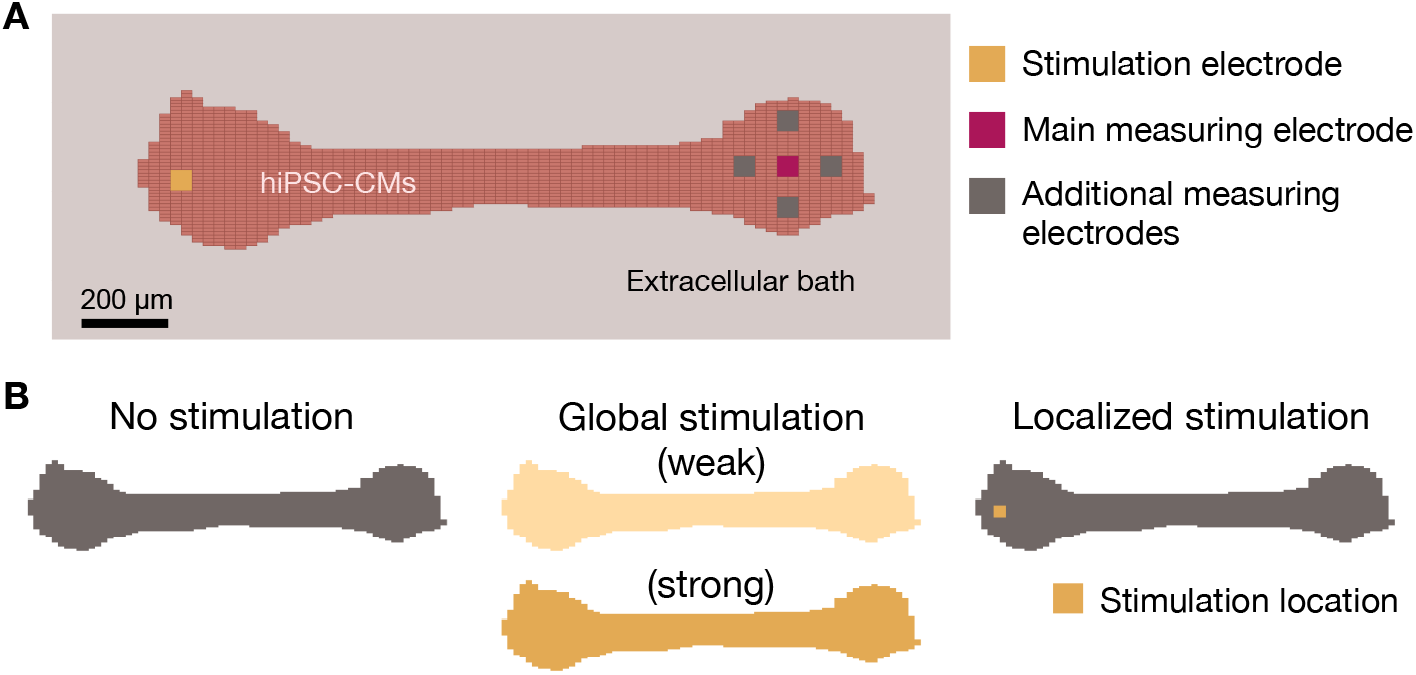
**A: Model geometry**. hiPSC-CMs are arranged in a dogbone-like geometry in a single well (see Chip C1 in [16]). Each cell has a length of 25 *µ*m and a width of 8 *µ*m. The location of one stimulation electrode and five measuring electrodes are indicated. Note that hiPSC-CMs are also located at the location of the electrodes. **B: Stimulation protocols**. We consider four different stimulation protocols: (i) no stimulation (spontaneous AP firing), (ii) weak global optical stimulation applied in the entire cell collection, (iii) same as (ii) but stronger stimulation, and (iv) localized extracellular stimulation through an electrode.

### 2.2 Stimulation approaches

In Figure 1B, we illustrate the stimulation approaches considered in this study. In the case of no stimulation, the cells beat spontaneously at a rate determined by their intrinsic pacemaker properties. The global stimulation is set up to mimic global optical pacing of the cells, represented by the activation of a negative 5 ms long transmembrane current. We consider two different strengths of this light activated current, one weak current of 3 *µ*A/cm^2^, which is barely enough to initiate an action potential and one strong current of 30 *µ*A/cm^2^, which is more than enough to initiate action potentials. For the localized stimulation, a 5 ms long constant current of 0.02 *µ*A is injected into the extracellular space through a stimulation electrode. This strength was determined to be strong enough to initiate local APs, but not so strong that it significantly affected the extracellular potential at the measuring electrodes. All stimulation is applied at 1 Hz.

### 2.3 Numerical methods

The KNM system is solved using a standard first order operator splitting scheme, splitting the non-linear single-cell dynamics from the linear part of the KNM system (see [15, 17, 18]) with a global time step of Δ*t* = 0.1 ms and a local time step of Δ*t* = 0.0005 ms for the forward Euler scheme used for the non-linear single-cell dynamics.

## 3 Results

### 3.1 Identical cells

We begin with a collection of identical, autonomously beating hiPSC-CMs using the computational model derived in [15, 16]. Figure 2 shows that cases (i), (ii) and (iii) generate no EPs even though all cases exhibit full transmembrane action potentials. In contrast, localized stimulation (iv) induces a traveling wave, leading to strong extracellular signals.

**Figure 2:**
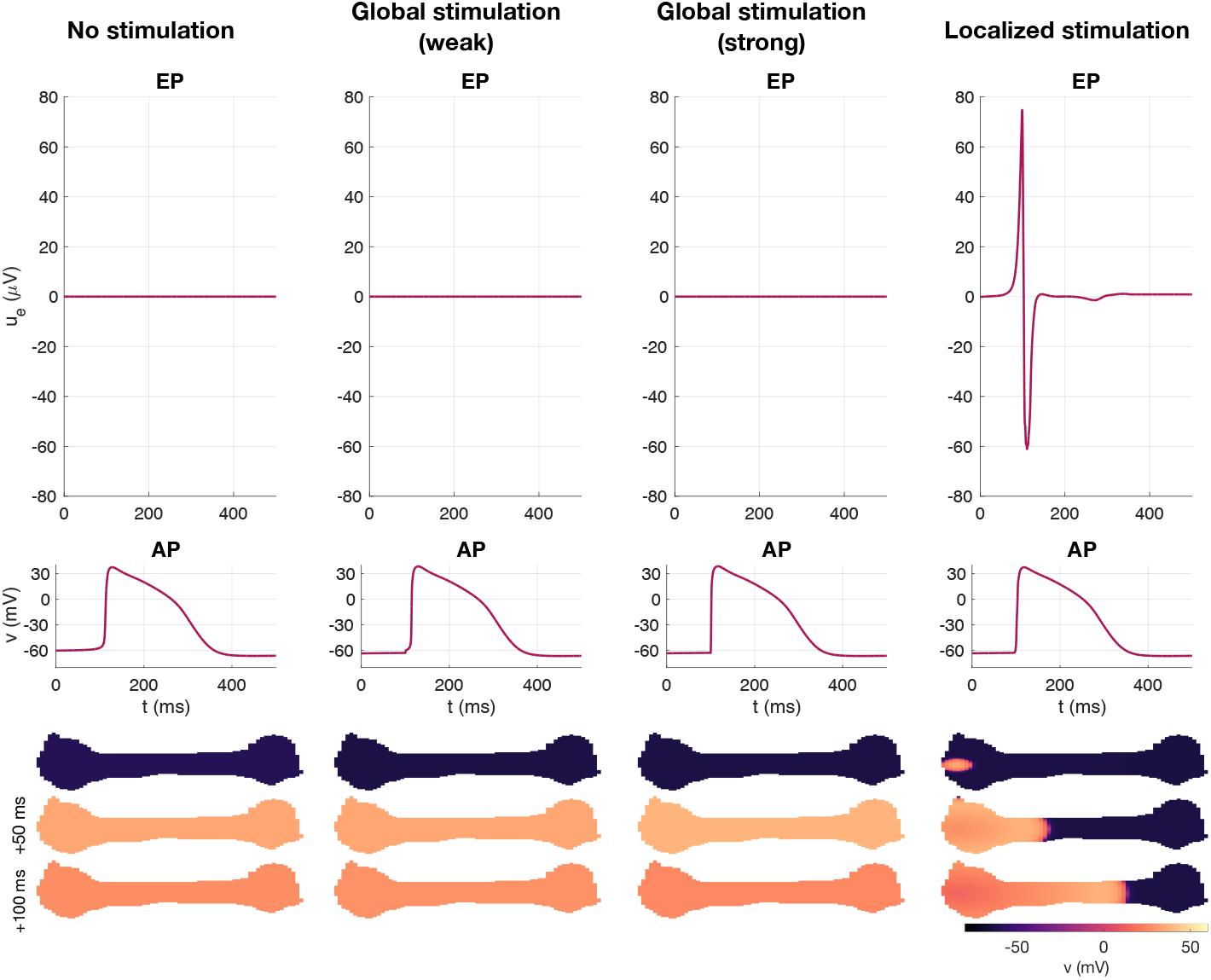
Identical cells. Simulation of a collection identical hiPSC-CMs for four different stimulation protocols. The upper panel displays the extracellular potential measured in the main measurement electrode illustrated in Figure 1A. The extracellular potential is zero in all cases except for localized stimulation. The next panel shows the membrane potential recorded at the location of the same electrode. All stimulation protocols display clear and similar action potential firing. In the lower panel, snapshots of the membrane potential are displayed for three points in time during AP firing. For the localized stimulation, an excitation wave is generated, whereas for the remaining stimulation protocols, the cells fire uniformly across the collection.

### 3.2 Varying cell properties

Figure 3 shows the effect of introducing cell-to-cell variation in the ion channel conductances. For each cell, a separate random scaling factor is drawn for each ionic current and used to modify the corresponding maximum conductance. The scaling factors are drawn from a normal distribution with expectation equal to the default conductance value and standard deviation of 20% of the default conductance value.

**Figure 3:**
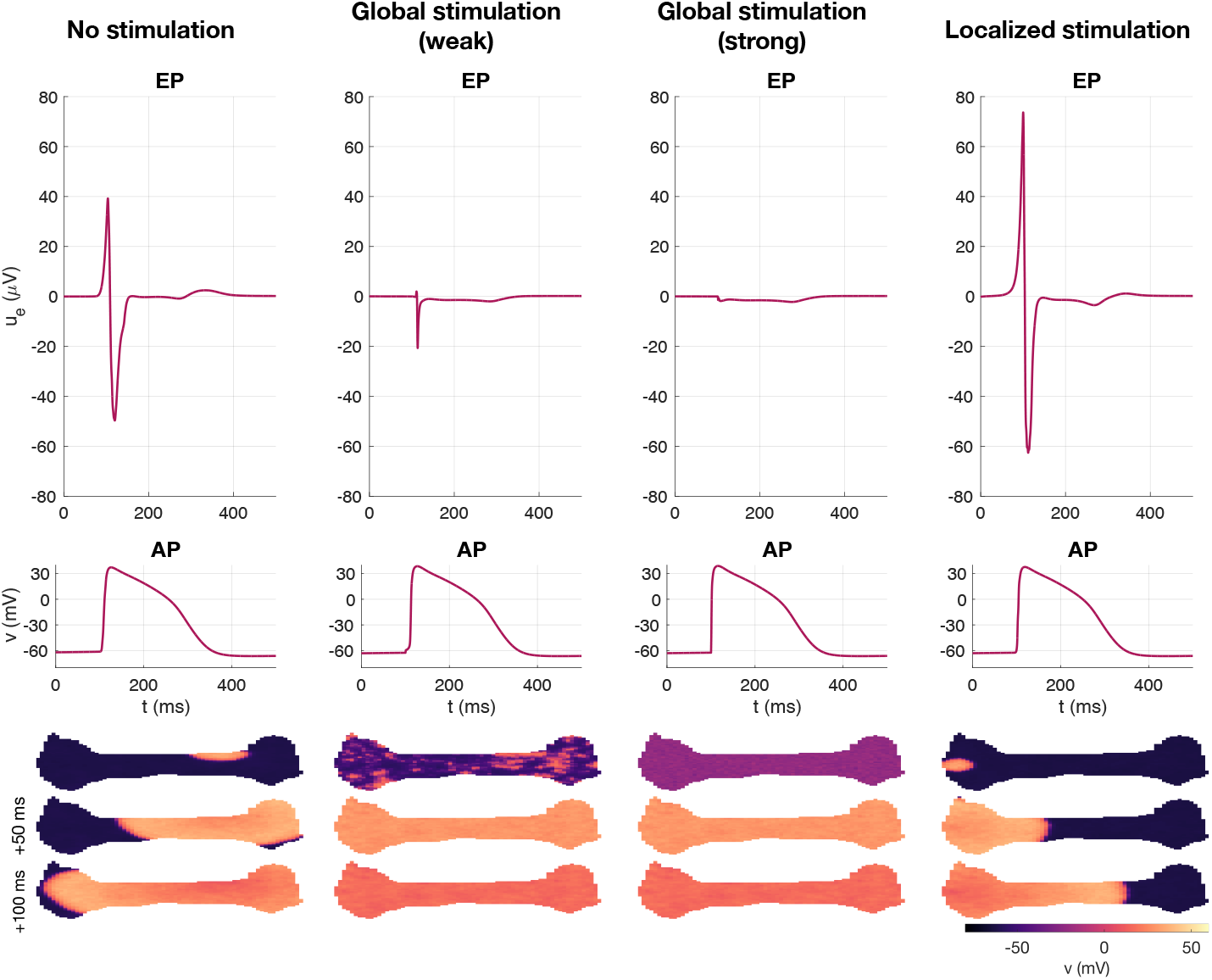
Varying cell properties. Simulations like in Figure 2, except that cell-to-cell variation is introduced in the conductances of the membrane currents (20% standard deviation). In this case, there are spatial gradients in the membrane potential for all stimulation protocols. Consequently, EPs are present in all cases, but wave initiation continues to yield the strongest signal.

In Figure 3, we observe that cell-to-cell heterogeneity enhances EPs in both the no stimulation (autonomous) case and in the global stimulation cases, but localized stimulation still produces the strongest and most reliable signal, driven by the wavefront. In the no stimulation case, the heterogeneous cell properties lead to action potential firing occurring at a location in the upper right part of the cell collection before the remaining cells fire, resulting in a traveling excitation wave similar to that generated in the localized stimulation case. For a weak global stimulation, certain spatial differences are present during depolarization, resulting in an extracellular signal that is visible, yet weaker than for the no stimulation and localized stimulation cases. For the strong global stimulation, the stimulation current evens out these spatial differences to a large extent, and the resulting extracellular signal is very weak.

### 3.3 Improved gap junction coupling

Due to the immature nature of hiPSC-CMs, gap junction coupling is not fully developed and may vary across different hiPSC-CM cell collections. For example, in [16], the conductance of the gap junctions connecting neighboring cells was found to vary between *G*_*g*_ = 1.7 nS and *G*_*g*_ = 180 nS for different cell collections. Figure 4 shows the effect of increasing the gap junction strength by a factor of 100 to *G*_*g*_ = 170 nS. We still consider 20% variation in cell properties. We observe that when the gap junction conductance is increased, the extracellular signal strength for the localized stimulation is considerably improved and consistently stronger compared to the other stimulation approaches.

**Figure 4:**
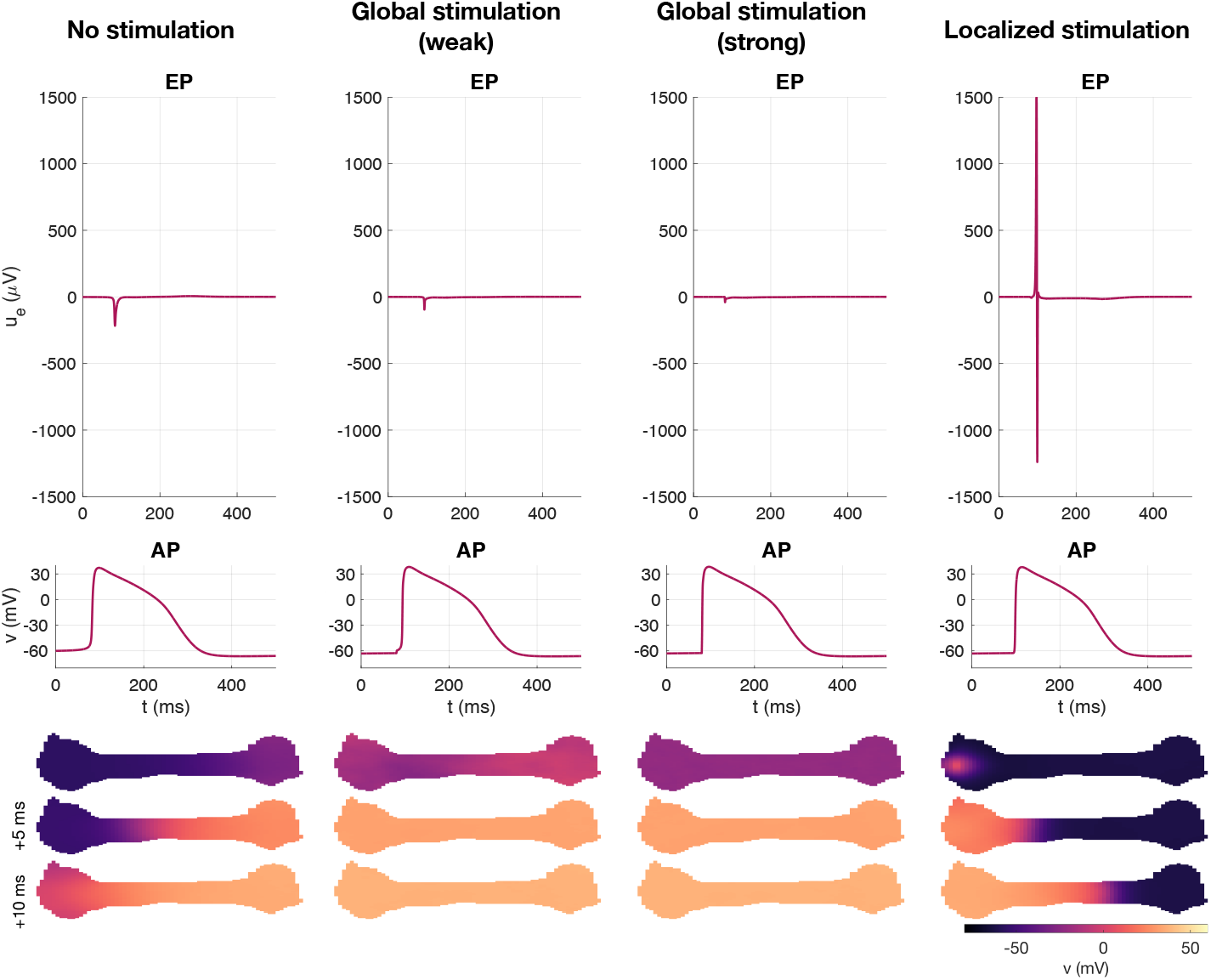
Improved gap junction coupling. Simulations like in Figure 2 except that the gap junction coupling strength is increased by a factor of 100, to *G*_*g*_ = 170 nS, and that there is 20% cell-to-cell variation in ion channel conductances. The increased gap junction coupling greatly enhances the EP signal strength for a localized stimulation.

### 3.4 Summary of perturbed properties

In Table 1, we have summarized how the maximum amplitude of the EP depends on different choices of the gap junction coupling strength and the degree of cell-to-cell variation. For all choices, the localized stimulated simulations yield the strongest EP signal. For any choice of gap junction coupling strength, 0% cell variation yields zero EP for the no stimulation and global stimulation cases. As cell variation is increased, the EP strength for the no simulation case and the global stimulation cases increases, but the localized stimulation yields the strongest EPs in all cases and in particular for strong gap junction coupling. In addition, the no stimulation case consistently give rise to stronger EPs than global stimulation, and a strong global stimulation results in the weakest EP signals of the considered approaches.

**Table 1:** Maximum amplitude of the extracellular potential (in *µ*V) over all five measurement electrodes illustrated in Figure 1. The cell-to-cell variation is represented by drawing ion channel conductances from a normal distribution with standard deviation of 0%, 5%, or 20%, and *G*_*g*_ represents the gap junction coupling strength between neighboring cells.

| Cell variation | $G_g$ | Stimulation | | | |
| --- | --- | --- | --- | --- | --- |
|  |  | none | global (weak) | global (strong) | localized |
| 0% | 1.7 nS | 0.0 | 0.0 | 0.0 | 74.8 |
|  | 17 nS | 0.0 | 0.0 | 0.0 | 468.8 |
|  | 170 nS | 0.0 | 0.0 | 0.0 | 1597.9 |
| 5% | 1.7 nS | 48.9 | 15.7 | 0.8 | 74.8 |
|  | 17 nS | 54.7 | 41.7 | 7.0 | 468.7 |
|  | 170 nS | 67.3 | 41.3 | 20.9 | 1607.2 |
| 20% | 1.7 nS | 49.7 | 35.0 | 3.3 | 73.7 |
|  | 17 nS | 144.0 | 140.2 | 28.0 | 465.3 |
|  | 170 nS | 232.9 | 176.9 | 83.5 | 1621.9 |

## 4 Discussion

The use of field potentials to assess drug-induced changes in hiPSC-CMs is well established [7, 19, 4, 5, 20, 21, 22, 23, 24]. Our results indicate that when analyses rely on extracellular potential (EP) recordings, the stimulation protocol plays a critical role in shaping signal strength. The clearest signals arise when stimulation initiates a traveling wave through the tissue. Autonomous activity without external stimulation can also yield measurable EPs, but uniform global stimulation consistently produces weaker signals. This effect is especially pronounced when strong global stimuli are used (see Table 1). Thus, if global stimulation must be used, the strength should be kept as low as possible while still reliably triggering action potentials, since stronger stimulation tends to suppress spatial voltage gradients and reduce EP amplitude.

When localized stimulation is applied, it must also be carefully tuned to avoid directly influencing the measurements; the observed signal should arise from the activity of the hiPSC-CMs, not from the input at the electrode. This is why the measurement electrodes are placed at the far end of the tissue in Figure 1. Furthermore, if the cells are too weakly coupled for supporting a traveling excitation wave, localized stimulation will fail to generate a wave, and relying on intrinsic beating is likely the best option. Finally, a further advantage of wave-based stimulation is that it permits measurement of the conduction velocity, which may provide an independent readout of sodium current strength [22, 25].

Variability in drug response detectability across hiPSC-CM platforms (Axion, Ncardia, Fujifilm, Pluricyte) has been noted in [26], particularly in settings where analysis relies on spontaneous activity. Our results suggest that part of this variability may arise from differences in how activation spreads across the tissue, not just from differences in cellular properties. In [27], extracellular signals remained below 2 *µ*V despite full action potentials, especially when activation was spatially uniform. This supports the idea that spatial voltage gradients – not just cellular excitability – are required for generating robust EPs. Enhanced spike amplitudes observed during electrical pacing in MEA studies [28] – often doubling the signal compared to spontaneous activity – are consistent with this interpretation. These findings may also help explain why uniform optogenetic stimulation sometimes fails to reveal subtle pharmacological effects [29].

## 5 Conclusion

The simulations of this report confirm that stimulation protocols have a strong influence on the strength of extracellular potentials (EPs) in small *in vitro* tissues. Stimulation that initiates a traveling wave produces spatial voltage gradients and robust EPs, while uniform global stimulation – despite its common use – yields much weaker signals. These results are directly relevant to the design of protocols for extracting electrophysiological biomarkers from hiPSC-derived cardiomyocytes.

## Notes

### Competing Interest Statement

Both authors are shareholders in Organos Inc., a start-up company specializing in measuring drug effects in microphysiological systems.
The authors hold the following patents/patent applications:
(a) US011846631B2 (2023) - Methods for determining drug effects on a mature cardiomyocyte. Tveito, A., Wall, S., and Jaeger, K.H. (granted),
(b) WO2021236535 (2021) - Determining drug effects using combination of measurements. Tveito, A., Jaeger, K.H., and Wall, S. (application being processed),
(c) WO2022264096 (2022) - Combination of existing drugs to repair the action potentials of cells. Jaeger, K.H., and Tveito, A. (application being processed).

